# RADAR: annotation and prioritization of variants in the post-transcriptional regulome of RNA-binding proteins

**DOI:** 10.1101/474072

**Authors:** Jing Zhang, Jason Liu, Donghoon Lee, Jo-Jo Feng, Lucas Lochovsky, Shaoke Lou, Michael Rutenberg-Schoenberg, Mark Gerstein

## Abstract

RNA-binding proteins (RBPs) play key roles in post-transcriptional regulation and disease. Their binding sites cover more of the genome than coding exons; nevertheless, most noncoding variant-prioritization methods only focus on transcriptional regulation. Here, we integrate the portfolio of ENCODE-RBP experiments to develop RADAR, a variant-scoring framework. RADAR uses conservation, RNA structure, network centrality, and motifs to provide an overall impact score. Then it further incorporates tissue-specific inputs to highlight disease-specific variants. Our results demonstrate RADAR can successfully pinpoint variants, both somatic and germline, associated with RBP-function dysregulation, that cannot be found by most current prioritization methods, for example variants affecting splicing.

## Background

Dysregulation of gene expression is a hallmark of many diseases, including cancer [1]. In recent years, the accumulation of transcription-level functional characterization data, such as transcriptional factor binding, chromatin accessibility, histone modification, and methylation, has brought great success to annotating and pinpointing deleterious variants. However, beyond transcriptional processing, genes also experience various delicately controlled steps, including the conversion of premature RNA to mature RNA, and then the transportation, translation, and degradation of RNA in the cell. Dysregulation in any one of these steps can alter the final fate of gene products and result in abnormal phenotypes[2-4]. Furthermore, the post-transcriptional regulome covers an even larger amount of the genome than coding exons and demonstrates significantly higher cross-population and cross-species conservation. Unfortunately, variant impact in the post-transcriptional regulome has been barely investigated, partially due to the lack of large-scale functional mapping.

RNA binding proteins (RBPs) have been reported to play essential roles in both co- and post-transcriptional regulation[5-7]. RBPs bind to thousands of genes in the cell through multiple processes, including splicing, cleavage and polyadenylation, editing, localization, stability, and translation[8-12]. Recently, scientists have made efforts to complete these post- or co-transcriptional regulome by synthesizing public RBP binding profiles[13-16], which have greatly expanded our understanding of RBP regulation. Since 2016, the Encyclopedia of DNA Elements (ENCODE) consortium started to release data from various types of assays on matched cell types to map the functional elements in post-transcriptional regulome. For instance, ENCODE has released large-scale enhanced crosslinking and immunoprecipitation (eCLIP) experiments for hundreds of RBPs[17]. This methodology provides high-quality RBP binding profiles with strict quality control and uniform peak calling to accurately catalog the RBP binding sites at a single nucleotide resolution. Simultaneously, ENCODE performed expression quantification by RNA-Seq after knocking down various RBPs. Finally, ENCODE has quantitatively assessed the context and structural binding specificity of many RBPs by Bind-n-Seq experiments[18].

In this study, we aimed to construct a comprehensive RBP regulome and a scoring framework to annotate and prioritize variants within it. We collected the full catalog of 318 eCLIP (for 112 RBPs), 76 Bind-n-Seq, and 472 RNA-Seq experiments after RBP knockdown from ENCODE to construct a comprehensive post-transcriptional regulome. By combining polymorphism data from large sequencing cohorts, like the 1,000 Genomes Project, we demonstrated that the RBP binding sites showed increased cross-population conservations in both coding and noncoding regions. This strongly indicates the purifying selection on the RBP regulome. Furthermore, we developed a scoring scheme, named RADAR (<u>***R***</u>N<u>***A***</u> Bin<u>***D***</u>ing Protein regulome <u>***A***</u>nnotation and p<u>***R***</u>ioritization), to investigate variant impact in such regions. RADAR first combines RBP binding, cross-species and cross-population conservation, network, and motif features with polymorphism data to quantify variant impact described by a universal score. Then, it allows tissue- or disease-specific inputs, such as patient expression, somatic mutation profiles, and gene rank list, to further highlight relevant variants (Fig. 1). By applying RADAR to both somatic and germline variants from disease genomes, we demonstrate that it can pinpoint disease-associated variants missed by other methods. In summary, RADAR provides an effective approach to analyze genetic variants in the RBP regulome, and can be leveraged to expand our understanding of post-transcriptional regulation. To this end, we have implemented the RADAR annotation and prioritization scheme into a software package for community use (radar.gersteinlab.org).

**Figure 1.**
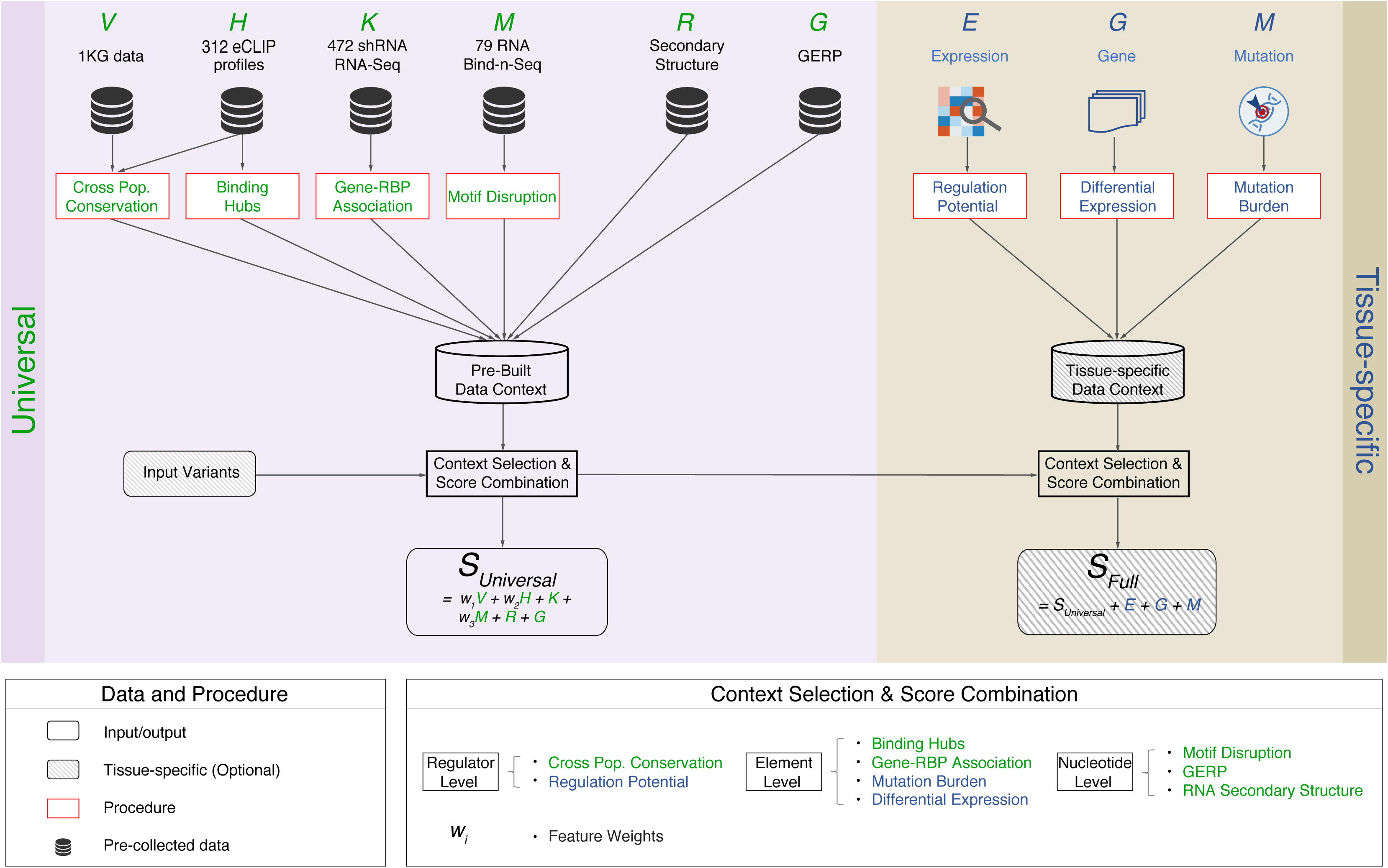
RADAR workflow. There are two RADAR score components: (1) a universal score derived from a pre-built data context including sequence and structural conservation, network centrality, motif, and knockdown information; (2) a tissue-specific (user-defined) score consisting of expression, gene, and mutation information to further highlight tissue-specific variants. The universal and tissue-specific score sum together to form the full RADAR score.

## Results

### Defining the RBP regulome using eCLIP data

We used the binding profiles of 112 distinct RBPs from ENCODE to fully explore the human RBP regulome (Additional file 1: Table S1), which has been previously under-investigated. Many of these RBPs are known to play key roles in post-transcriptional regulation, including splicing, RNA localization, transportation, decay, and translation (Additional file 2: Figure S1 and Table S1).

Our definition of the RBP regulome covers 52.6 Mbp of the human genome after duplicate and blacklist removal (Fig. 2A). It is 1.5 and 5.9 times the size of the whole exome and lincRNAs, respectively. In addition, only 53.1% of the RBP regulome has transcription-level annotations, such as transcription factor binding, open chromatin, and active enhancers. 55.1% of the RBP regulome is in the immediate neighborhood of the exome regions, such as coding exons, 3’ or 5’ untranslated regions (UTRs), and nearby introns (Fig. 2C; see methods section and Additional file 2: Table S2 for more details). Furthermore, we observed significantly higher cross-species conservation score in the peak regions versus the non-peak regions in almost all annotation categories, providing additional evidence of regulatory roles of RBPs (Fig. 2C). In summary, the large size of the regulome, the limited overlap with existing annotations, and the elevated conservation level highlight the necessity of computational efforts to annotate and prioritize the RBP regulome.

**Figure 2.**
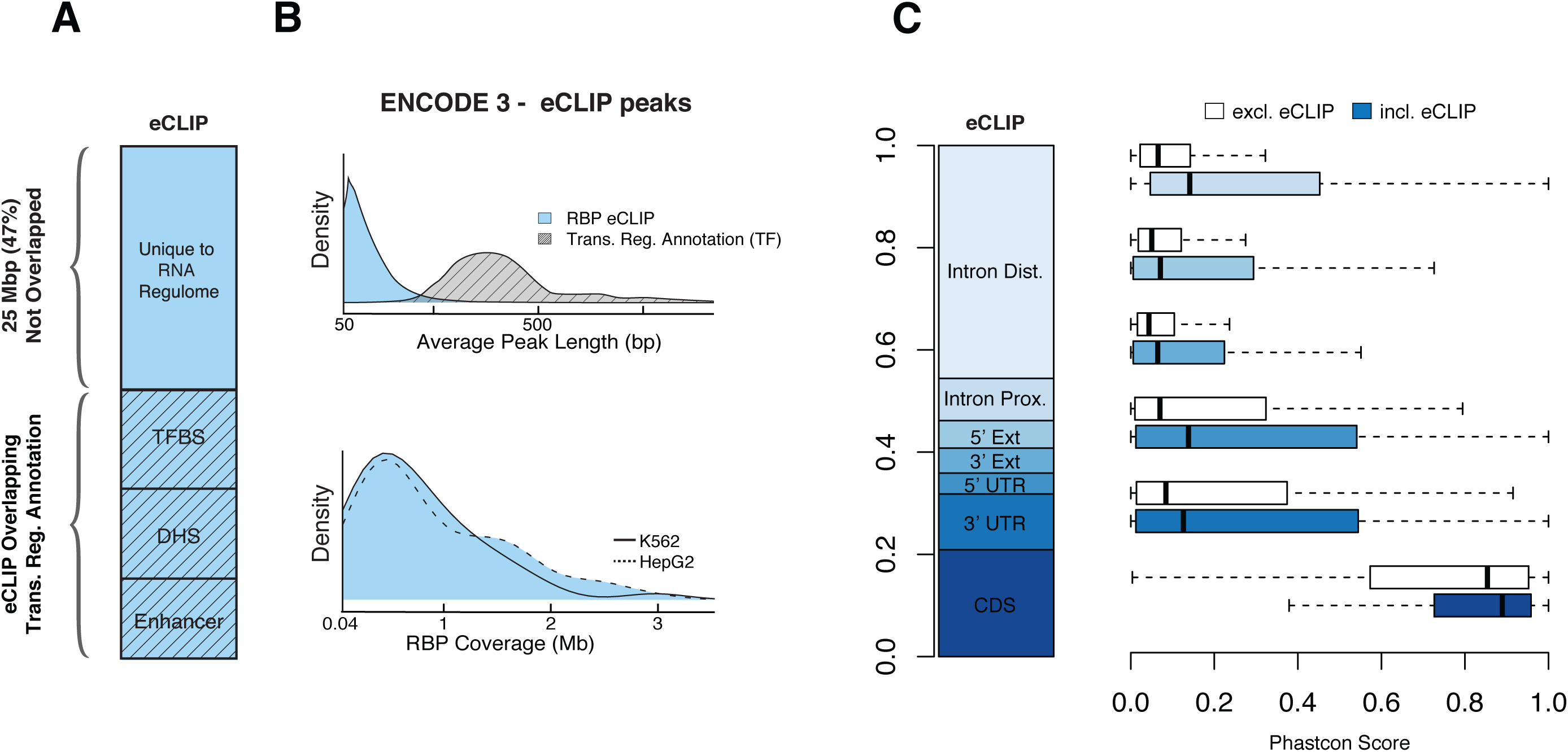
RBP regulome and cross-species conservation. (A) Intersection of eCLIP peaks versus transcriptional level annotations, with 25Mbp unique to the RBP regulome; (B) Average length of binding peak for RBP eCLIP data versus TF ChIP-Seq and the similar distribution of RBP coverage between K562 and HepG2 cell lines; (C) Fraction of RBPs falling into each annotation category as well as boxplots of PhastCons scores of annotations intersecting peaks (blue) versus annotations with no intersections (white).

### Using universal features for RADAR score

To annotate and prioritize variants in RBP binding sites, we built a universal score framework for RADAR that includes three components: (1) sequence and structure conservation; (2) network centrality; and (3) nucleotide impact from motif analysis.

#### Sequence and structure conservation in the RBP regulome

Cross-species sequence comparisons have been widely used to discover regions with biological functions [19, 20]. For example, GERP score maps the human genome to other species to identify nucleotide-level evolutional constraints[21, 22]. Therefore, we used the GERP score in our RADAR universal framework to detect potentially deleterious mutations in the RBP regulome (see methods section for more details).

Since the enrichment of rare variants indicates a purifying selection in functional regions in the human genome[19, 23, 24], here we inferred the conservation of RBP binding sites by integrating population-level polymorphism data from large cohorts (i.e. the 1,000 Genomes Project)[25, 26]. GC percentage may confound such inference by introducing read coverage variations, which is a sensitive parameter in the downstream variant calling process[27, 28]. Therefore, we calculated the fraction of rare variants, defined as those with derived allele frequencies (DAFs) less than 0.5%, within the binding sites of each RBP. Then we compared them with those from regions with similar GC content as a background (see methods section for more details). In total, 88.4% of the RBPs (99 out of 112) showed elevated rare variant fraction in coding regions after GC correction (Fig. 3A). Similarly, in the noncoding part of the binding sites, 93.8% of RBPs (105 out of 112) exhibited an enrichment of rare variants. This observation convincingly demonstrates the accuracy of our RBP regulome definition (Additional file 1: Table S2).

**Figure 3.**
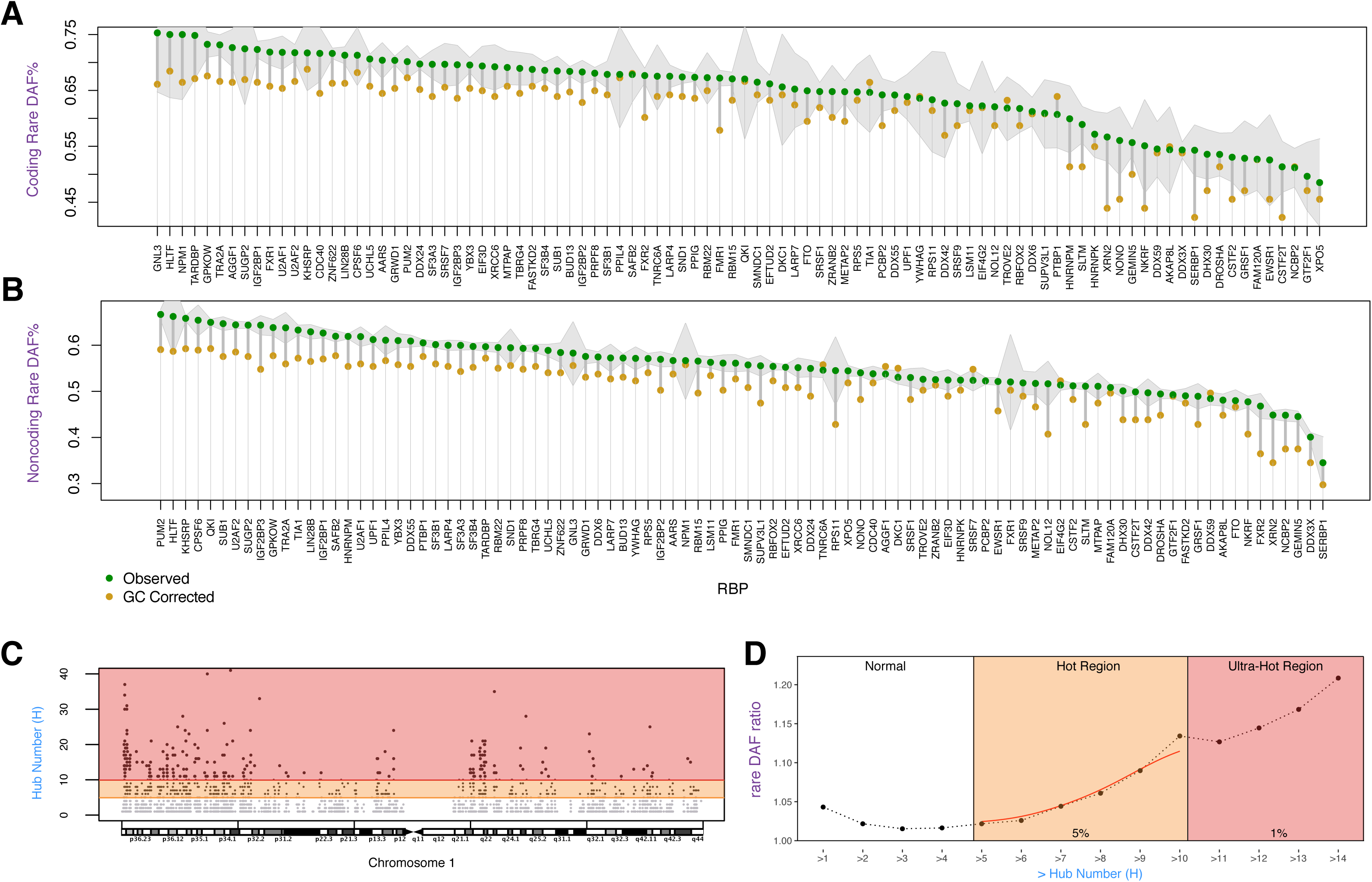
Cross-population conservation of RBP peaks and binding hubs. (A-B) Rare variant percentage in coding/noncoding regions. The green dot represents RBP peaks, and yellow dot represents the genome average after GC correction. Shaded lines are the 95% confidence interval of the rare variant percentage of the RBP peaks; (C) an example of RBP binding hubs. Red and orange shadings denote regions with the top 1% (ultra-hot) and 5% (hot) RBPs binding; (D) corrected rare variant percentage at positions with different cumulative hub numbers.

Some well-known disease-causing RBPs demonstrate the largest enrichment of rare variants. For example, the oncogene XRN2, which binds to the 3’ end of transcripts to degrade aberrantly transcribed isoforms, showed significant enrichment of rare variants in its binding sites[29]. Specifically, it demonstrates 12.7% and 10.3% more rare variants in coding and noncoding regions, respectively (adjusted P values at 1.89×10^-9^ and 2.85×10^-118^ for one-sided binomial tests)[30]. Hence, we used the enrichment of rare variants to infer the selection pressure in RBP binding sites and adjust the universal variant scores in such regulator regions (see methods for more details).

RNA secondary structures have been reported to affect almost every step of protein expression and RNA stability[31]. We incorporated structural features predicted by Evofold, which uses a phylogenetic stochastic context-free grammar to identify functional RNAs in the human genome that are deeply conserved across species[32]. We found that the RBP binding sites demonstrated significantly higher conversation after intersecting with conserved structural regions defined by Evofold (Additional file 2: Figure S2). Thus, we used the Evofold regions in our universal scoring system.

#### Highlighting variants in binding hubs

It has been reported that genes within network hubs demonstrate higher cross-population conservation — a sign of strong purifying selection[23, 24, 33]. We hypothesized that RBP binding hubs could show similar characteristics because once mutated they might introduce larger regulation alterations. To test this, we separated the regulome based on the number of associated RBPs. Most regulome regions (62%) were associated with only one RBP (Additional file 2: Figure S3). As the number of RBPs increased, we observed a clear trend of larger rare variant enrichment (Fig. 3D). For instance, noncoding regions with at least five or 10 RBPs exhibited 2.2% or 13.4% more rare variants, respectively (top 5% and 1%, Fig 3D). This observation supports our hypothesis that the RNA regulome hubs are under stronger selection pressure and, therefore, should be given higher priority when evaluating functional impact of mutations (details in Additional file 2: Figure S4, S5, and S6).

#### Emphasizing genes differentially expressed after RBP knockdown

RNA-seq expression profiling before and after shRNA mediated RBP depletion from ENCODE can help to infer the gene expression changes introduced by RBP knockdown. Variants with disruptive effects on RBP binding may affect or even completely remove the RBP binding and hence affect gene expressions in a similar way. Therefore, we extracted the differentially expressed genes from RNA-Seq before and after shRNA-mediated RBP depletion (Additional file 1: Table S3). Then, we up-weighted all variants that were located near the differentially expressed genes (Additional file 1: Table S4) and simultaneously disrupted the binding of the corresponding RBPs (schematic in Additional file 2: Figure S7).

#### Using motif analysis to determine nucleotide level impact

Mutations that change the RBP binding affinity may alter RBP regulation via motif disruption. We quantified the difference of position weight matrix (PWM) scores of the mutant allele against the reference allele. RADAR consists of two sources of motifs. First, we used the motifs identified from RNA Bind-n-Seq experiments from ENCODE because it has been reported that many RBP binding events *in vivo* can be captured by binding preferences *in vitro*. Second, we used the *de novo* motifs discovered directly from binding peaks using the default settings in DREME (see details in methods). For each variant, we quantified the nucleotide effect using the highest motif score from these two sources.

### Incorporating user-specific features to reweight variant impact

Variant prioritization can be improved if informative priors can be appropriately incorporated into the scoring system. Therefore, our RADAR framework allows various types of user-inputs to help identify disease-relevant variants. Specifically, we adopted a top-down scheme to incorporate regulator and element level information to up-weight factors that are possibly associated with disease of interest.

#### Highlighting key regulators through expression profiles

Key regulators are often associated with disease progression, so variants that affect such regulation should be prioritized[34]. RADAR finds such key regulators by combining the RBP regulatory network information with expression profiles. Specifically, for cancer, we first constructed the RBP network from eCLIP binding peaks and used the TCGA data to define the gene differential expression status from disease and normal cell types (see Methods). Then for each RBP, we quantified its regulation potential by associating its network connectivity with aggregated disease-to-normal differential expressions from many samples using regression. We applied this approach on 19 cancer types from TCGA and the regulation potentials are given in Fig. 4. The values of the regulation potential (*β*_1_, see Methods) for all cancer types and RBPs are provided in Additional file 1: Table S5). We found that among the RBPs with larger regulation potential, many have been reported as cacer-associated genes (Additional file 1: Table S6). Our regression approach was also performed on a patient level, and survival analysis based on each patient’s regulatory potential was performed (see Fig. 4C). Interestingly, the regulatory potential of two key RBPs PPIL4 and SUB1 were found to be significantly associated with patient survival (Fig. 4C). In our RADAR framework, we further highlight variants that are associated with RBPs with high regulation potential in their corresponding cancer types by adding extra score to their disease-specific scores (see more details in methods). We can easily extend such analysis for other diseases by incorporating differential expression profiles from others cohorts such as GTEx[35, 36].

**Figure 4.**
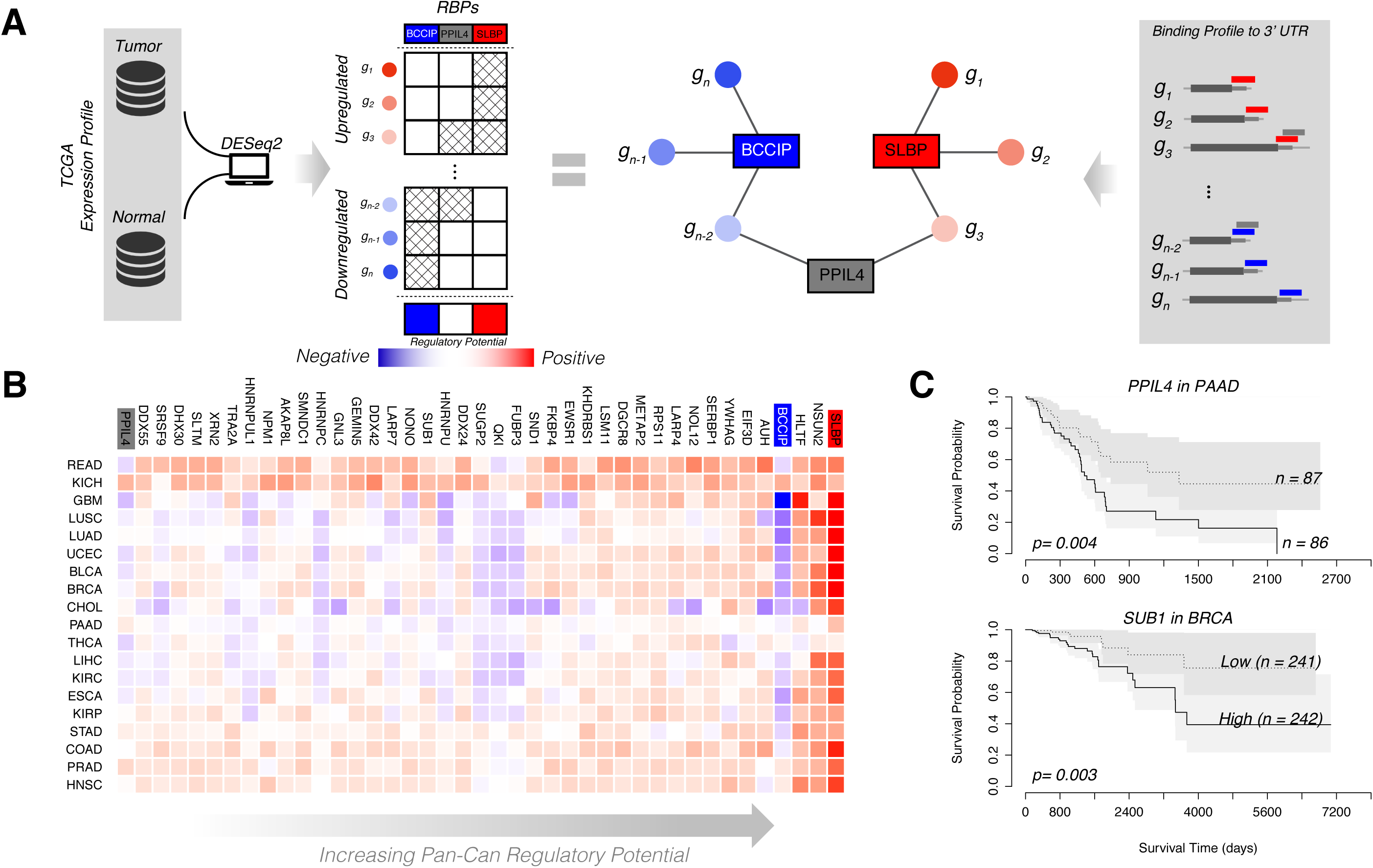
Regulation potential inference of RBPs. (A) Schematic of RBP regulation potential calculation; (B) Heatmap of RBP regulation potential in 19 cancer types; (C) RBPs associated with patient survival. Patient survival data from TCGA, survival analysis performed using *R* package *survival* (2.42-3). Differential expression within a patient is calculated as the difference between tumor and normal expression, converted to a Z-score.

#### Up-weighting key elements from either prior knowledge or mutational profiles

RADAR reconsiders the functional impact difference among RBP peaks by their associated genes. For example, genes that undergo significant expression or epigenetic changes are mostly cell-type-specific and can be used to highlight more relevant variants. Currently, our RADAR framework can up-weight all the RBP peaks that are close to genes with significant differential expression (DESeq2[37]).

In addition, RADAR can incorporate somatic variant recurrence, which has been widely used to discover key disease regions, to reweight different RBP peaks. Peaks with more somatic mutations than expected are often considered to be disease-driving[38-40]. Here, we first defined a local background somatic mutation rate from a large cohort of cancer patients to evaluate the mutation burden in each RBP peak (see details in methods). Variants that are associated with burdened elements are given higher priority in our scoring scheme.

### Prioritizing variants with a RADAR weighted scoring scheme

By integrating the pre-built and user-specific data context described above, our scoring scheme evaluated the functional impacts of variants that are specific to post-transcriptional regulation (Fig. 1 and Table 1). We used an entropy-based criterion to up-weight rarer annotations. First, RADAR added up the (universal) score of variants for all pre-built features, which include sequence and structure conservation, network binding hub, RBP-gene association, and motif information, using the entropy-based weights. Then, depending on user inputs, RADAR further up-weighted variants’ score based on tissue specific information from mutations in the key RBP binding sites, nearby genes with differential expression, or the RBP regulatory potential (See Additional file 2: Figure S8 for more details).

**Table 1.** Features used by RADAR

| Category | Feature | Source | Scoring Scheme |
| --- | --- | --- | --- |
| Universal | Cross-population conservation | eCLIP | Adjusted-entropy |
|  | Cross-species conservation | Gerp | Sigmoid Function |
|  | Structural conservation | EvoFold | Fixed Value |
|  | RBP Binding hub | eCLIP | Adjusted-entropy |
|  | RBP-gene association | shRNA RNA-seq | Fixed Value |
|  | Motif disruption | Bind-n-Seq<br>DREME | Adjusted-entropy |
| User-specific | RBP regulatory potential | Expression | Fixed Value |
|  | Differentially expressed Genes | Prior knowledge | Fixed Value |
|  | Mutation Recurrence | Mutation profiles | Fixed Value |

### Applying RADAR to pathological germline variants

We calculated the universal RADAR scores on all pathological variants from HGMD (version 2015) and compared them with variant scores from 1,000 Genomes variants as a background. As expected, the HGMD variants are scored significantly higher (Additional file 2: Figure S9). For example, the mean RADAR score for HGMD variants is 0.589, while it is only 0.025 for 1,000 genomes variants (P-value <2.2×10^-16^ for one-sided Wilcoxon test). We further compared the universal RADAR scores of HGMD and 1,000 genomes variants within the RBP regulome to remove potential bias since HGMD variants may be more likely to be within or nearby to exons. We still observed significantly higher universal RADAR score in the HGMD ones (1.871 vs. 1.337, P-value <2.2×10^-16^ for one-sided Wilcoxon test, Additional file 2: Figure S9).

We further compared RADAR scores of HGMD variants with other methods. In total, 720 HGMD variants were explained by our methods but could not be highlighted by other methods (see details in methods, Additional file 1: Table S7). Many of these variants are located nearby the splice junctions. An example is shown in Fig. 5A (more details in Additional file 1: Table S8). This variant is located 4 base pairs away from splice junction in BRCA1. eCLIP experiments showed strong binding evidence in 5 RBPs (Fig. 5A). Specifically, the T to C mutation strongly disrupts the binding motif of PRPF8, increasing the possibility of splicing alteration effects. Our finding is not highlighted in previous methods for variant prioritization, such as FunSeq, CADD, and FATHMM-MKL (Additional file 2: Table S3).

**Figure 5.**
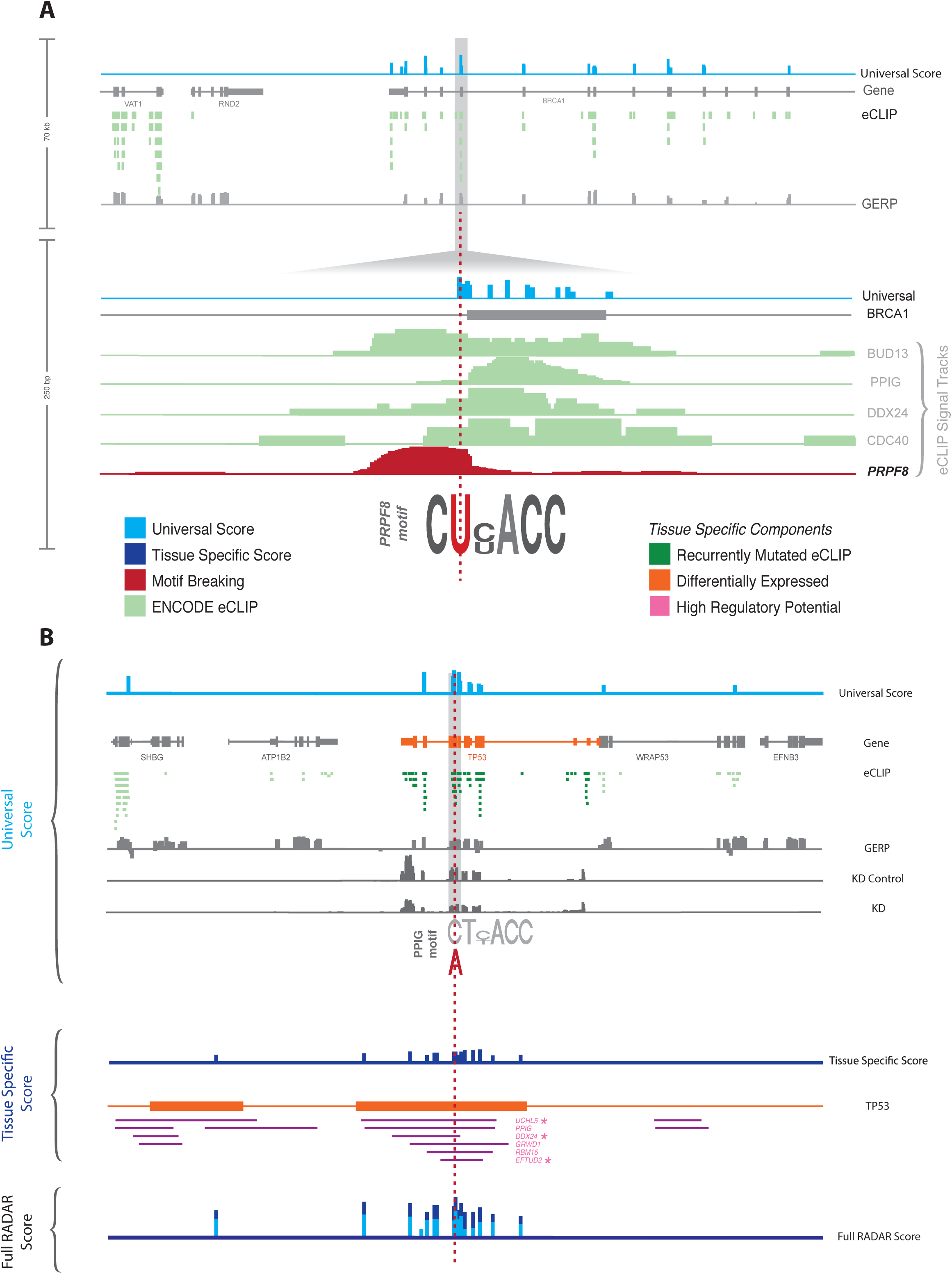
Example of breast cancer somatic variant with high overall RADAR score. (A) An example of a top scoring BRCA1 intron HGMD variant highlighted by the universal RADAR score. This variant is in an RBP binding hotspot, shows high GERP score, and breaks the motif of the splicing factor PRPF8 (red); (B) We selected an exonic variant with high RADAR score on chromosome 17 as an example. It was inside an RBP binding hub with a high GERP score and breaks the motif of PPIG. It also has several tissue-specific features, like within the well-known cancer-associated gene TP53 (orange track) and its associated binding peaks were significantly burdened (dark green). Also, adding expression profiles from TCGA shows that 3 out of the 6 RBPs binding there demonstrated high regulation potentials in driving tumor-specific expression pattern. All these external pieces of evidence further boost this variant’s tissue-specific score.

### Applying RADAR universal score to somatic variants in cancer

We next aimed to leverage our universal RADAR scheme to evaluate the deleteriousness of somatic variants from public datasets. Due to the lack of a gold standard, we evaluated our results from two perspectives. First, we reasoned that since hundreds of cancer-associated genes are known to play essential roles through various pathways[41, 42], variants associated with these genes are likely to have the higher functional impact[23]. To test this hypothesis, we first selected variants within the 1kb region of the COSMIC[43] genes and compared them with other variants. We tested four cancer types, breast, liver, lung, and prostate cancer, and found in all cases that variants associated with COSMIC genes showed significant enrichment, with a larger RNA level functional impact (Fig. 6 and Additional file 2: Figure S10). For example, we found a 4.58- and 8.75-fold increase in high-impact variants at a threshold level of 3 and 4, respectively, in breast cancer patients (P < 2.2×10^-16^, one-sided Wilcoxon test).

**Figure 6.**
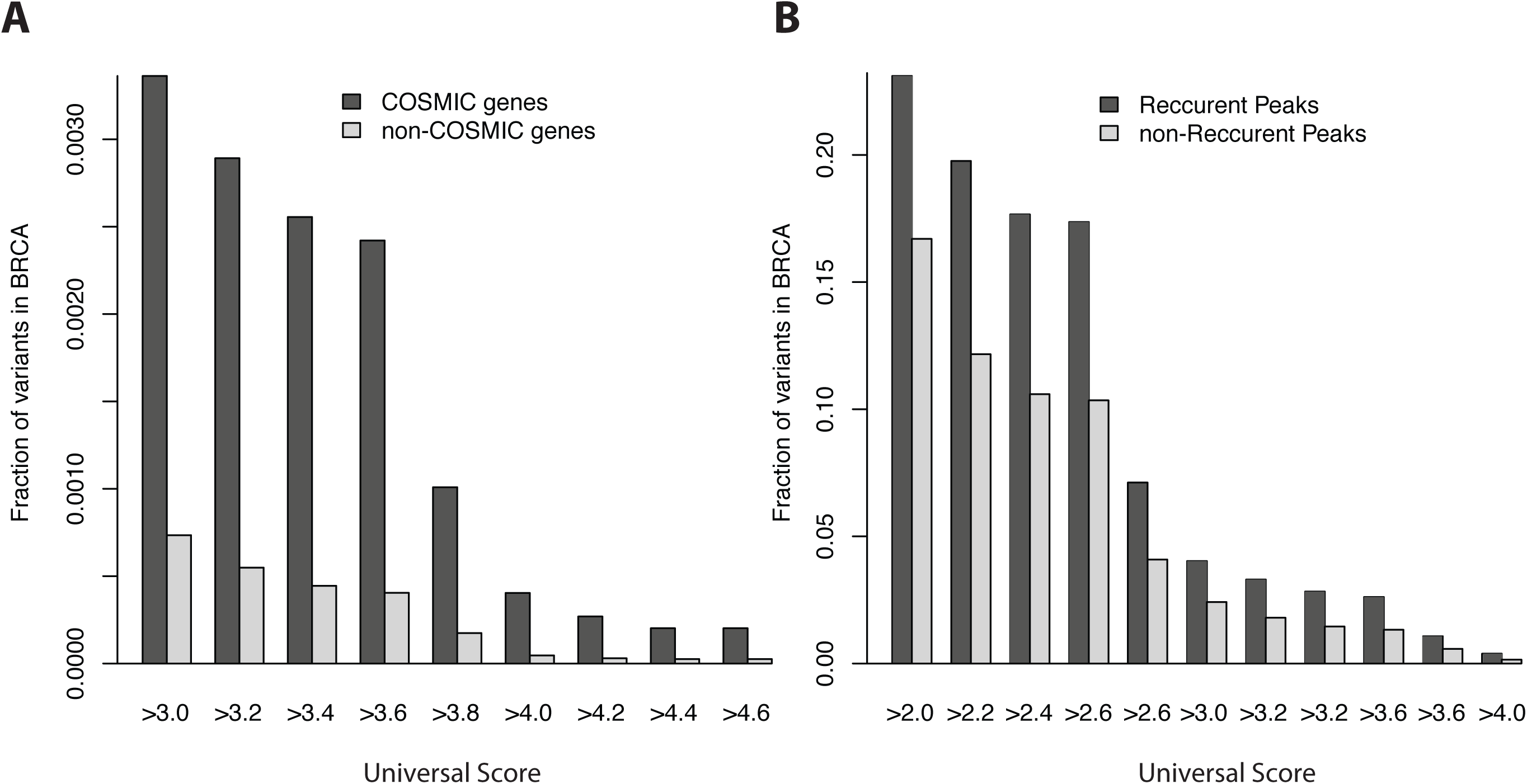
Performance of universal RADAR score on somatic variants. (A) Enrichment of high RADAR universal score variants associated with COSMIC genes in breast cancer; (B) Enrichment of high RADAR universal score variants within RBP peaks with recurrent variants in breast cancer.

In our second approach, we hypothesized that variant recurrence could be a sign of functionality and may indicate an association with cancer[19, 23, 24]. Thus, we compared the variants’ scores from RBP binding peaks with or without recurrence. Specifically, we separated the RBP peaks with variants mutated in more than one sample from those that were mutated in only one sample, and then compared the universal RADAR scores. We found that in most cancer types, peaks with recurrent variants were associated with a larger fraction of high-impact mutations. For example, in breast cancer recurrent elements demonstrated a 1.67-, and 2.57-fold more high-impact variants with RADAR greater than 3.0 and 4.0, respectively, resulting in a P value of 2.2×10^-16^ in one-sided Wilcoxon test. We observed a similar trend in most of the other cancer types (Additional file 2: Figure S10).

### A case study on breast cancer patients using disease-specific features

We applied our method to a set of breast cancer somatic variants from 963 patients released by Alexandrov *et al.* [44]. We used COSMIC gene list, expression and mutational profiles as additional features. In total, we found that around 3% of the 687,517 variants could alter post-transcriptional regulation to some degree. We incorporated the above disease-specific features and demonstrated how they could help to reweight the variant scoring process on a coding variant in Fig. 5B. This variant is located within an RBP binding ultra-hot region and showed high sequence conservations (7% more rare variants for its binding RBP). It also demonstrated strong motif disruption effect (PPIG in Fig 5B). All such features resulted in a universal RADAR score of 3.67, which is ranked 290 out of all variants. However, we found that it is located in the exon region of the well-known tumor suppressor TP53 (orange track in Fig.5B), and its binding peaks demonstrated more than expected somatic mutations (purple in Fig. 5). Besides, 3 out of the 6 RBPs binding there showed high regulation potential in breast cancer (green in Fig. 5). Hence, these additional features boost its overall RADAR score to 6.67, which is ranked 47 out of all variants). In comparison, this variant only shows moderate scores for FunSeq2 (3) and CADD (7.46), and while it is scored in the top but showed much lower rank than RADAR.

RADAR aims to prioritize variants relevant to the post-transcriptional regulome, while FunSeq2, FATHMM-MKL, and CADD focus on those that affect the transcriptional regulome. Therefore, we do find many variants that demonstrate a high overall RADAR score, but only show moderate FunSeq2, CADD, and FATHMM-MKL scores. For example, 127 coding and 78 noncoding variants that are ranked within the top 1% of overall RADAR scores are not in the top 10% of CADD, FunSeq2, or FATHMM-MKL scores (Additional file 1: Table S9 and T10). Many of such variants are located in RBP binding hubs, and undergo strong purifying selection, demonstrated strong motif disruptiveness, and are regulated by key RBPs that are associated with breast cancer from multiple sources of evidence (Additional file 2: Figure S11). We believe the discovery of such events demonstrates the value of RADAR as an important and necessary complement to the existing transcriptional-level function annotation and prioritization tools.

## Discussion

In this study, we integrated the full catalog of eCLIP, Bind-n-Seq, and shRNA RNA-Seq experiments from ENCODE to build an RNA regulome for post-transcriptional regulation. Our defined RBP regulome is remarkably larger than one may think. It covers up to 52.6 Mbp of the genome (Fig. 2A) and the majority of it is not covered by previous annotations focusing on transcriptional-level regulation, such as DHS, TFBS, and enhancers. We found that the RBP regulome demonstrated noticeably higher conservation in two aspects: higher cross-species conservation in almost all annotation categories (Fig. 2C) and higher across-population conservation by showing significant enrichment in rare variants (Fig. 3). These two sources of evidence support the notion that the RBP regulome is under strong purifying selection and carries out essential biological functions. Also, these results signify the necessity of computational tools to annotate and prioritize variants in the RBP regulome. Furthermore, we showed that the cross-population conservation in eCLIP binding sites demonstrates strong importance when predicting disease versus common germline variants (Additional file 2: Figure S12 and Table S4).

By integrating a variety of regulator-, element-, and nucleotide-level features, we propose an entropy-based scoring frame, RADAR, to investigate the impact of somatic and germline variants. The variant prioritization framework of RADAR contains two parts. First, by incorporating eCLIP, Bind-n-Seq, shRNA RNA-seq experiments with conservation and structural features, we built a pre-defined data context to quantify the universal variant impact score. This approach is suitable for multiple-disease analysis or cases where no other prior information can be used. We applied this RADAR universal score to HGMD pathological variants and highlighted many candidates that cannot be highlighted by other methods. Besides, our RADAR framework provided detailed explanations of the underlying disease-causing mechanism (Fig. 5A). In addition to the universal score, RADAR also allows user-specific inputs such as prior gene knowledge, patient expression and mutation profiles for a re-weighting process to highlight relevant variants in a disease-specific manner. As an example, we applied the RADAR disease-specific scores to variants from several cancer types and showed that RADAR could identify relevant variants in key cancer-associated genes (Fig. 5B). We additionally showed that the RADAR framework trained in a closely matched cell type has better power to pinpoint pathological variants in a particular disease, which is promising as new eCLIP experiments are performed in various other cell types (see Additional file 2: Figure S13 and S14).

It is important to note that as compared to ChIP-Seq experiments which generate peaks with up to kbp resolution, eCLIP experiments provide higher resolution functional site annotation (even single nucleotide resolution). Such accurate and compact annotation can greatly improve our variant function interpretations. We also want to mention that most of the current eCLIP peak calling approaches call peaks on the annotated transcribed regions (Additional file 2: Figure S15). With the development of computational approached for eCLIP peak calling, we hope that the size of our annotated RBP regulome can be further expanded.

## Conclusions

In summary, we have shown that RADAR is a useful tool for annotating and prioritizing post-transcriptional regulome for RBPs, which has not been covered by most of the current variant impact interpretation tools. Our method provides additional layers of information to the current gene regulome. Importantly, the RADAR scoring scheme can be used in conjunction with existing transcriptional-level variant impact evaluation tools, such as FunSeq [23, 24], to quantify variant impacts. Given the fast-expanding collection of RBP binding profiles from additional cell types, we envision that our RADAR framework can better tackle the functional consequence of mutations from both somatic and germline genomes.

## Methods

### eCLIP Data Processing and Quality Control

We collected 318 eCLIP experiments of 112 unique RBPs from the ENCODE data portal (encodeprojects.org, released and processed by July 2017). eCLIP data was processed through the ENCODE 3 uniform data processing pipeline and peaks with score 1,000 were used in our analysis. We then removed peaks overlapped with blacklisted regions. We further separated the peaks into coding regions and the noncoding regions in our analysis to infer the selection pressure. We also provide versions of the eCLIP peaks that are annotated by RBP’s function, such as splicing – which is the most common function aside from RNA binding (see radar.gersteinlab.org).

### Universal RADAR Score

#### Cross-population conservation inference

The cross-population conservation score consists of two components. The Shannon entropy considers the length effect of the RBPs while the selection pressure inference aims to determine the conservation of regions. For the Shannon entropy, for each RBP, we define *f* to be

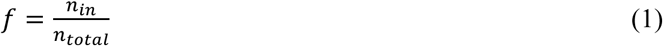

where *n*_*in*_ represents the number of 1,000 Genomes variants falling in that RBP peaks, and *n*_*total*_ to be the total number of 1,000 Genomes variants (fixed number). In this way, *f* considers the binding site coverage of an RBP, since a larger coverage is more likely to have a larger value of *n*_*in*_. The Shannon entropy is therefore equal to

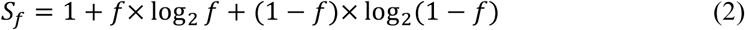

We then calculate the selection pressure from the enrichment of rare germline variants from the 1,000 Genomes Project. Our analysis at each step is separated into coding and noncoding parts. For a given RBP, we suppose its binding peaks contain *n*_*r*_ rare variants (*DAF* ≤ 0.005) and *n*_F_ common variants. The percentage of rare variants in that RBP’s binding peaks is defined as

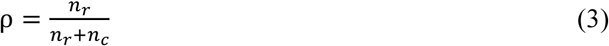

The value of ρ is often confounded by factors such as GC content. To correct for potential GC content bias, we bin the genome into 500 base pair bins and group them according to their GC percentage. Then we compute the background rare variant percentage using the same rare and common variants from 1,000 Genomes Project for each group (see Additional file 2: Figure S16 and S17). For a given RBP with GC percentage *g*, we select the background group with closest GC, to obtain a background rare variant percentage 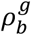. Therefore, after adjusting for GC bias, the enrichment of rare variants is defined to be

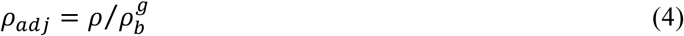

RBPs with a *ρ*_*adj*_ larger than 1 suggests a higher than expected selection pressure. We then adjust the population conservation entropy score as follows

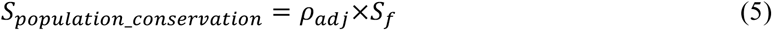

Given a variant falling in the RBP regulome that intersects a set of RBP eCLIP peaks, set P, the cross-population conservation score of that variant is equal to the maximum *S*_*population_conservation*_ for all RBPs in set P.

#### Cross-species conservation using GERP

We use GERP score to measure the cross-species conservation. For each position, a GERP of greater than 2 is often used to define bases that are conserved. The transformation of GERP to a RADAR component score, is adapted from Fu *et al*. Therefore, a sigmoidal transformation is used to fit the GERP scores between 0 and 1, and the parameters used force the curve to be sharp at GERP equal to 2 (Additional file 2: Section 13.2).

#### Structural Conservation

We use the output of Evofold as an indicator of cross species RNA structure. A variant falling in a region given by Evofold as conservative receives a score of 1 while a variant that does not fall in such region receives a score of 0.

#### RBP Binding Hubs and Networks

We define the number of RBPs binding at a position to as *H*. We first separated the RBP peaks into coding and noncoding regions and then grouped regions on the genome based on *H*. For each group of regions with hub number *H*, we calculated the GC-corrected enrichment of rare variants *ρ*_*adj | H*_ for each group in coding and noncoding regions by equation (3, 4). We determine the hub numbers, *H*_*normal*_, *H*_*hot*_, and *H*_*ultra_hot*_ associated with normal, hot, and ultra-hot regions, respectively. *H*_*hot*_ and *H*_*ultra_hot*_ are associated with the top 5% and 1% of binding RBPs (Figure 3) to represent rare and ultra-rare events. Our values of *ρ*_*adj* | *H*_ are altered in such a way to reflect this phenomenon. The 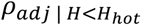 associated with hub scores less than that of the hot regions are converted to 0. The 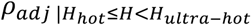 associated with hot regions demonstrate a mostly increasing trend and are smoothed using a kernel smoother. The 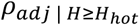 of the ultra-hot region is kept constant, equal to the max 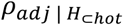 of the hot region.

We also compute *f* for the set of regions associated with a fixed *H* (From equation 1). For each value of *f*, we compute the Shannon entropy from equation (2) to be *S*_*H*_. Finally, we multiply the values of the *ρ*_*adj*_ by its respective Shannon entropy, S. A variant falling in the regulome with hub score *H* would have a score equal to the *ρ*_*adj* | *H*_ * *S*_*H*_.

#### Motif Analysis and Disruption

We used the changes of PWMs introduced by a variant to quantify the motif disruptiveness effect through motiftools (https://github.com/gersteinlab/MotifTools). Specifically, we defined the disruption score, *D*_*score*_, as in equation (6) to represent the difference between sequence specificities to an alternative sequence.

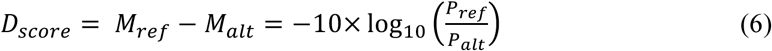

where *P*_*ref*_ and *P*_*alt*_ are the PWM scores from the reference and alternative allele. Here, the motif scores for reference and alternate sequences are given as

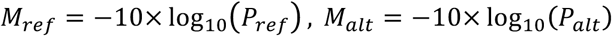

To quantify a motif breaking event, we require that the *p*-value for the reference allele is at least 5×10^−4^. There are two motif sources in our analysis. First, we identified RBP motifs using DREME software (Version 4.12.0) directly from RBP peaks[45]. Then, we also incorporated motifs from RNA Bind-N-Seq (RBNS) [18] to characterize sequence and structural specificities of RBPs. For each variant that affected multiple RBP binding profiles, we used the max score (Additional file 2: Table S5). A threshold of *D*_*score*_ > 3 is used to describe a disruption event that is significant, and a variant having a *D*_*score*_ less than this threshold receives a score of 0. For variants receiving a *D*_*score*_ larger than the threshold, we additionally compute the Shannon entropy given for a variant with *D*_*score*_ = *D* as

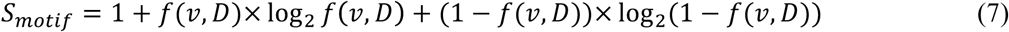

where *f(*v, *D*) represents the number of 1,000 Genomes Project variants, *v*, that have a *D*_*score*_ greater than *D* divided by the total number of 1,000 Genomes Project variants. This entropy is then weighted by the population conservation score.

#### RBP-gene association using shRNA RNA-Seq

To determine if an RBP, *R*, is associated with a gene, *g*, we intersect the peaks of *R*, with the transcript annotation of *g*. If an intersect exists, we form a linkage between the intersected peak of *R* and *g*. If some variant falls in that specific peak of *R*, the variant significantly disrupts the motif of RBP *R*, and gene *g* demonstrates at least a 2.5-fold change in its expression after KD of RBP *R*, we give the variant an additional score of 1.

### Tissue Specific Score

#### RBP Regulatory Potential

RADAR allows inputs in addition to the pre-built context to calculate the disease-specific variant score. In this paper, we used the TCGA expression profiles as an example on the cancer variant prioritization. Specifically, we downloaded expression profiles of 19 cancer patients of 24 types from TCGA. In order to get a robust differential expression analysis, we excluded several cancer types that have less than 10 normal expression profiles and used DESeq2 [37] to find tumor-to-normal differentially expressed genes (corrected P from DESeq2 <0.05). Let 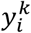 represent the differential expression status of gene *i* of the *k*^*th*^ cancer type.

We inferred the regulatory power of each RBP, R, through a regression approach of the above differential expression and RBP network connectivity as

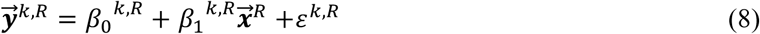

where 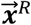 is the binary connectivity vector for all genes and *R* (1 if the gene is a target, else 0). We used the absolute value of *α*_1_^*k,R*^ to indicate the regulation potential of RBP *R* in cancer type *k*. If a variant falls in a region with at least one RBP binding, and at least one of the *p*-values associated with *α*_1_^*k,R*^ is significant, then we consider variants falling in that particular RBP to have an additional score of 1.

#### Recurrence in Somatic Mutations

We prioritized variants in RBP binding sites are with more-than-expected somatic mutations. To evaluate the somatic mutation burden, we first separated the genome into 1Mbp bins and calculated a local background mutation rate in each window. Then for each eCLIP peak, we counted the number of somatic mutations, and compared it to the nearest local 1Mbp context using a one-sided binomial test. If a specific RBP binding site was enriched for somatic mutations, the variant falling in that site was given a higher priority.

#### Differentially Expressed Key Genes

For each peak of each RBP, we find the associated gene of that peak by intersecting with the Gencode gene definitions. Using the DESeq2 results, we consider genes with q-values that are less than 0.05 to be differentially expressed. If an RBP peak is associated with a gene that is significantly differentially expressed in a tissue type, we increase the score of the variant falling in such peak by 1.

## List of abbreviations

RBP: RNA binding Protein;
ENCODE: the Encyclopedia of DNA Elements;
PWM: position weight matrix;
eCLIP: enhanced crosslinking and immunoprecipitation;
RADAR: <u>***R***</u>N<u>**A**</u> Bin<u>***D***</u>ing Protein regulome <u>***A***</u>nnotation and p<u>***R***</u>ioritization;
UTR: untranslated regions;
DAF: derived allele frequency;
RBNS: RNA Bind-N-Seq.

## Declarations

### Ethics approval and consent to participate

Not applicable.

### Consent for Publication

Not applicable.

### Availability of data and materials

We have made this RNA variant annotation and prioritization tool available as an open-source software at https://github.com/gersteinlab/RADAR[46]. The source code is released under MIT License. The website contains details on usage, examples, resources, and dependencies. We also provided a genome-wide pre-built RADAR baseline score for every base pair on the genome (hg19 and GRCh38 version of the genome). We released all pre-built data context necessary for RADAR and these genome-wide baseline scores at Zenodo which can be found at https://doi.org/10.5281/zenodo.1451838 [47]. Users can also directly query the annotation and functional impact score from radar.gersteinlab.org. See more details in Additional file 2: Section 15.

The eCLIP and shRNA datasets used in RADAR are from the ENCODE project and available at encodeproject.org[48]. We used germline single nucleotide polymorphisms from the 1000 Genomes Project, Phase 1[26]. Somatic variants are collected from Alexandrov *et al*[49]. The GERP score is from[21, 22].

The RNA-seq expression data and clinical survival data is from the TCGA project[50]. Gene annotations are from the GENCODE Project[51, 52].

### Authors’ contributions

JZ, JL and MG conceived the study and wrote the manuscript. JZ, JL, DL, LL, JF wrote the framework and performed the method evaluation. DL and JF developed the website. LL and MR carried out studies associating RNA structure. JL, DL, and SL participated in motif analysis. All authors have read and approved the final manuscript.

## Acknowledgements

We thank Brenton Graveley, Gene Yeo, Peter Freese, and Eric Van Nostrand for useful discussion on eCLIP and RBNS data.

## Additional Files

Additional file 1: Large Supplemental Tables from RADAR providing results and summaries regarding the data used, but are too large to be put in PDF format. (XLSX, 2.2 MB)

Additional file 2: Supplemental Figures and Tables for RADAR providing more information on results and methods. (PDF, 3.6 MB)

